# Covariation analysis with improved parameters reveals conservation in lncRNA structures

**DOI:** 10.1101/364109

**Authors:** Rafael C A Tavares, Anna Marie Pyle, Srinivas Somarowthu

**Affiliations:** Department of Chemistry, Yale University, New Haven, Connecticut, USA; Department of Molecular, Cellular and Developmental Biology, Yale University, New Haven, Connecticut, USA; Howard Hughes Medical Institute, Chevy Chase, Maryland, USA; Department of Biochemistry and Molecular Biology, Drexel University College of Medicine, Philadelphia, Pennsylvania, USA

## Abstract

The existence of phylogenetic covariation in base-pairing is strong evidence for functional elements of RNA structure, although available tools for identifying covariation are limited. R-scape is a recently developed program for prediction of covariation from sequence alignments, but it has limited utility on long RNAs, especially those of eukaryotic origin. Here we show that R-scape can be adapted for powerful prediction of covariation in long RNA molecules, including mammalian lncRNAs.

---

Long non-coding RNAs (lncRNAs) are well accepted as crucial regulators of gene expression and disease progression^1^. Despite the ubiquity and significance of lncRNAs, our understanding of structure-function relationships within this class of molecules is extremely limited^2^. Studies of ribozymes, riboswitches, viral RNAs, mRNA UTRs and even coding sequences have shown that conserved RNA secondary and tertiary structures are vital for RNA function^3, 4^ It has therefore been of interest to determine whether lncRNA molecules contain regions of functional structure and whether these structures are conserved^5-7^. If conservation in base-pairing could be established, it would provide powerful evidence that RNA structure plays a role in aspects of lncRNA function. Several empirical studies have demonstrated the existence of structured regions within lncRNAs, and conventional phylogenetic covariation analyses were found to support the empirically-determined structures^8-10^. Indeed in at least two cases, these modules of RNA structure were flanked by highly conserved sequences that are consistent with a biological role for lncRNA substructures^9, 10^.

However, a powerful new method for stringent determination of nucleotide covariation, known as R-scape, failed to support the existence of conserved base-pairings in well-studied functional lncRNAs such as Xist and HOTAIR^11^. On the basis of these findings, it was concluded that those lncRNAs do not contain conserved structure and are therefore unlikely to contain functional elements of discrete structure. Like many tools for phylogenetic analysis, R-scape was developed for application to small, highly structured RNA molecules for which many sequences are available (such as bacterial riboswitches). We reasoned that, at least in its current form, R-scape might not be equipped to confront the challenges posed by large, multidomain eukaryotic RNA molecules. We therefore set out to test the limitations of R-scape covariation analysis and to determine whether the approach could actually be modified in order to successfully identify conservation of structures in mammalian lncRNAs.

A major challenge for the analysis of eukaryotic lncRNAs is the severe limitation in available sequences^12^. We reasoned that this limitation, rather than any inherent lack of evidence for lncRNA structure, might explain the reported inability of R-scape to identify conserved structure in mammalian lncRNAs. To test this hypothesis, we analyzed the ability of R-scape to detect basepair covariation in seven well-characterized, highly structured RNA molecules (tRNA, 5S ribosomal RNA, 5.8S ribosomal RNA, eukaryotic RNase P, U2 snRNA, U5 snRNA and the eukaryotic small subunit ribosomal RNA) using input alignments that were restricted in three different ways: 1) Inclusion of the original RFAM seed alignment 2) Sub-sampled alignments and 3) Restriction to mammalian sequences. In the sub-sampled RFAM alignments, we limited the number of sequences and the average pairwise identity to control for effects arising solely from restrictions in these parameters (See Methods, Figure 1). The alignments restricted to mammalian sequences represent the currently available alignments that have been built for most lncRNAs. Not surprisingly, there is a precipitous drop in covariation support for most of these test RNAs in both the ‘sub-sampled’ and ‘mammalian sequence’ conditions. Eukaryotic RNase P (> 300 nt) is the most dramatic example, as only 13% of the base pairs can be flagged as covariant by R-scape in the sub-sampling analysis (Figure 1). It is also worth highlighting the particular case of 5.8S rRNA, for which the RFAM seed alignment already has a relatively high pairwise sequence identity (~68%). Predictably, R-scape finds covariation support for only 44% of the base pairs in the 5.8S rRNA structure, and no support (0%) upon restriction of the analysis to mammalian sequences. In fact, with the exception of tRNA, for which even mammalian sequences have high nucleotide diversity, R-scape was unable to detect the majority of covarying base pairings in these model RNAs when the input alignments were limited to mammals. These results indicate that R-scape fails to detect covariation not just in lncRNAs, but in most of the structurally complex, well-characterized functional RNA molecules that have been tested.

**Figure 1.**
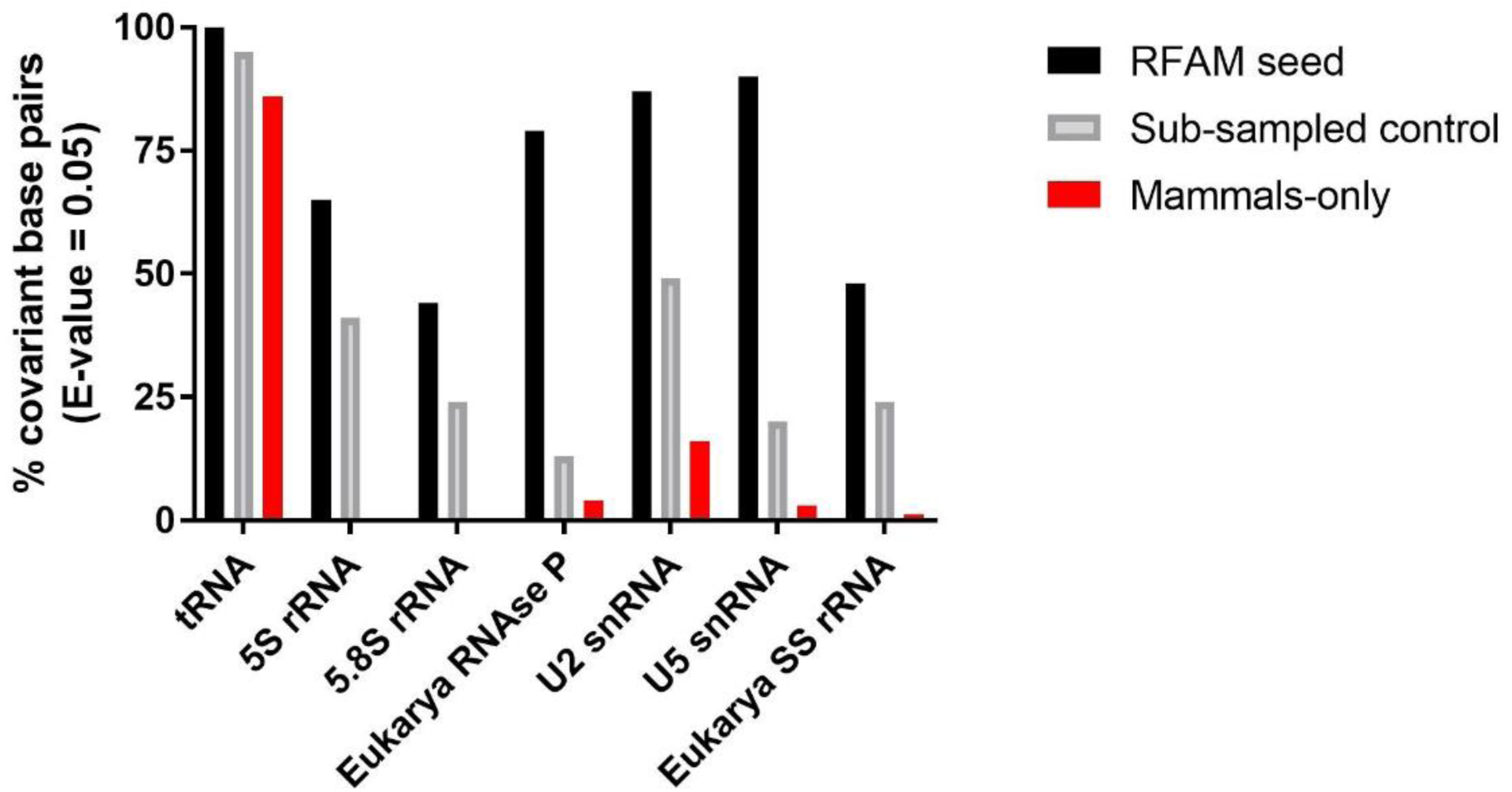
Restriction in alignment characteristics (number of sequences, average pairwise sequence identity and phylogenetic diversity) significantly impair R-scape’s ability to detect covariation in highly conserved structured RNAs. The percentage of covariant basepairs flagged by R-scape is shown in the graph for each tested RNA alignment.

It is important to note that RFAM alignments are hand-curated and refined^13^, therefore, deviations from RFAM’s ideal heuristics may bias R-scape results. This phenomenon was shown to be true for other covariance prediction algorithms when RFAM alignments were compared to emulated genomic alignments as inputs^14^. Multiple sequence-based alignments from datasets like the TBA/Multiz (UCSC genome browser) can be used to build covariation models and generate structural alignments for lncRNAs, but these alignments lack the quality of RFAM alignments, which can then affect R-scape prediction sensitivity. Finally, since genomic alignments may not accurately reflect the regions of lncRNA loci that are actively expressed, there is a consistent need for direct characterization and annotation of lncRNA transcripts across species in order to improve identification of conserved sequence and structure motifs, as described elsewhere^15, 16^.

There is accumulating evidence that lncRNAs possess local modules of RNA structure and that they can contain both structured and unstructured regions^8, 10^. Given that R-scape uses the entire length of an RNA sequence for analysis, it is possible that the presence of unstructured regions negatively impacts the ability of R-scape to identify structural conservation. To test this, we analyzed the ability of R-scape to predict covariation when unstructured regions are included in an alignment. R-scape is reported to perform well on riboswitches, using sequences that are restricted to the functional, structured region of the molecule. We therefore chose the SAM-I riboswitch (RF00162) as an example, but we now included the surrounding mRNA regions from the alignment. The mRNA regions were aligned using MAFFT^17^, and the alignment for the SAM-I riboswitch region was kept the same as in the RFAM alignment. We then compared R-scape predictions by varying the number of sequences in the alignment. In the case of the SAM-I riboswitch alone, R-scape predicted significant covariation even with only 40 sequences in the alignment (Figure 2A), as reported previously. However, inclusion of the flanking mRNA in the alignment resulted in a notable decrease in R-scape performance: Even when 60 sequences are included in the alignment, R-scape could identify covarying base pairs in only one helix (Figure 2B), indicating that the presence of unstructured RNA regions has a strong influence on R-scape analysis output. However, as the number of sequences in the alignment increases, R-scape can identify more covarying base pairs, even when unstructured regions are included. This suggests that R-scape may ultimately become a powerful tool for identifying covarying base-pairs when a sufficient number of sequences are provided (> 90 for SAM-I riboswitch). However, since the alignments for most human lncRNAs are currently limited to 30-60 mammalian sequences, R-scape default settings should not be applied to lncRNA covariation analysis.

**Figure 2.**
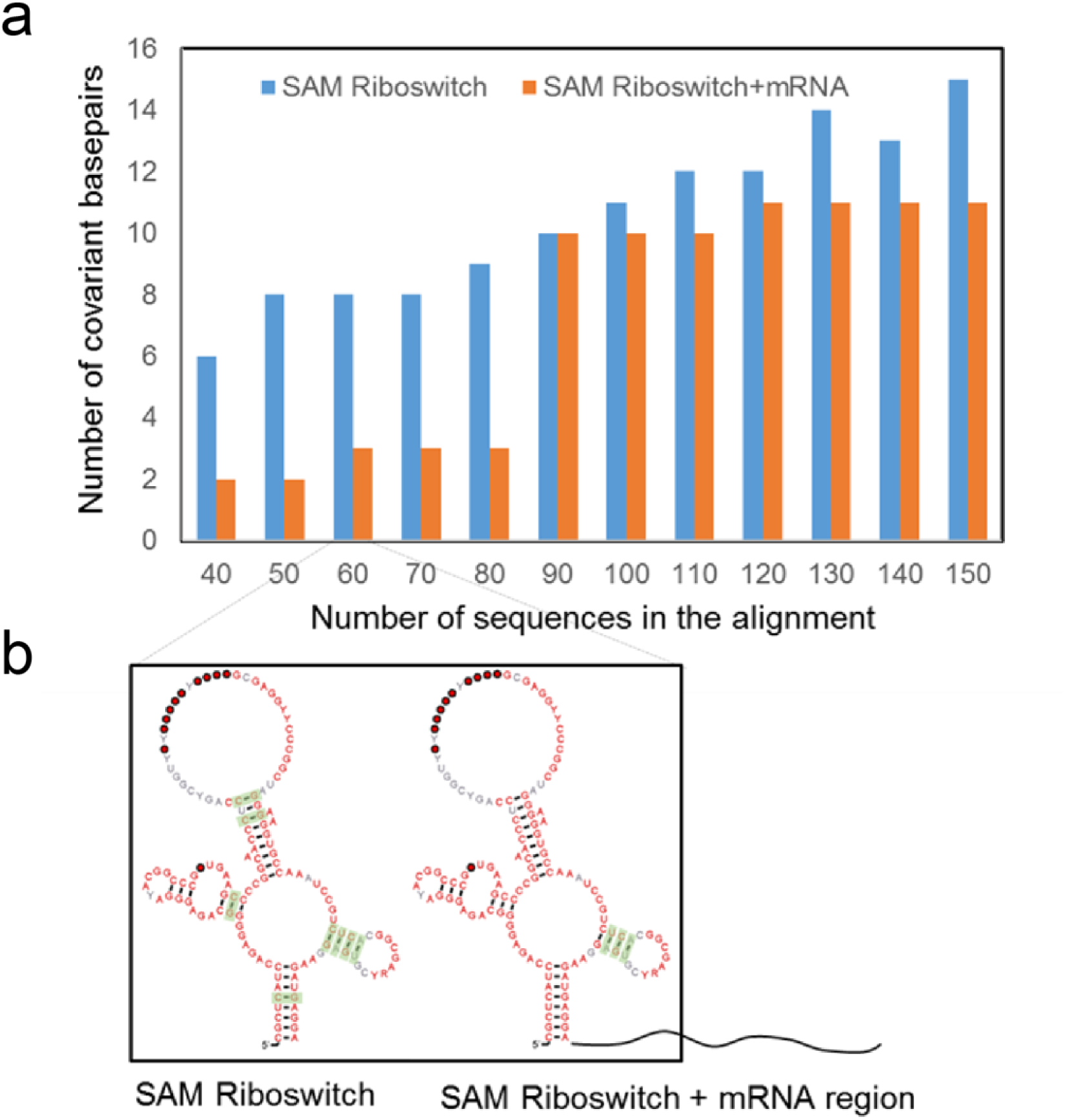
R-scape analysis on the SAM-I riboswitch (RF00162) with and without unstructured mRNA regions in the alignment. (**a**) Sensitivity of R-scape to the presence of adjacent unstructured regions, as a function of the number of sequences in the alignment. (**b**) Influence of an adjacent unstructured region on predicted covariation in the SAM-I riboswitch, using 60 sequences in the alignment. The figure shows the graphical output of each analysis generated by R-scape using R2R drawing notation. Green boxes indicate covariant basepairs. Consensus nucleotide letters are colored according to their sequence conservation in the alignment as given by percent identity thresholds (75% identity in gray; 90% identity in black; 97% identity in red). Individual nucleotides are represented in circles according to their positional conservation in the alignment corresponding to percent occupancy thresholds (50% occupancy in white; 75% occupancy in gray; 90% occupancy in black; 97% occupancy in red).

Another feature that is expected to influence the performance of any covariation analysis is the length of an RNA molecule and of its corresponding structural alignment. LncRNAs are typically very large and many exceed 1kb^18^. However, R-scape was benchmarked with a test set consisting predominantly of small RNAs. Of the 104 RNAs in that test set^11^, there are only 21 RNAs with an average length greater than 200 nts and only seven that exceed 1kb, and all the seven are ribosomal RNAs. It is therefore unlikely that the R-scape default parameters are appropriate for analysis of large RNAs. To test this, we asked whether R-scape performs better when the analysis is broken down in short overlapping windows tiling the entire RNA rather than when given a long whole-length alignment. We examined alignments (see methods) of two long RNAs in sliding windows: 1) 7SK RNA (RFAM ID:00100) and 2) Aphthovirus internal ribosome entry site (RFAM ID: 00210). For both RNAs, R-scape was able to identify more covarying base-pairs when the analysis was run with sliding windows than when given the full-length alignment (Fig. 3), indicating that the R-scape default parameters work better on short alignments, either as aligned sequences of inherently small RNAs or long RNA alignments that have been analyzed in a set of sliding windows.

**Figure 3.**
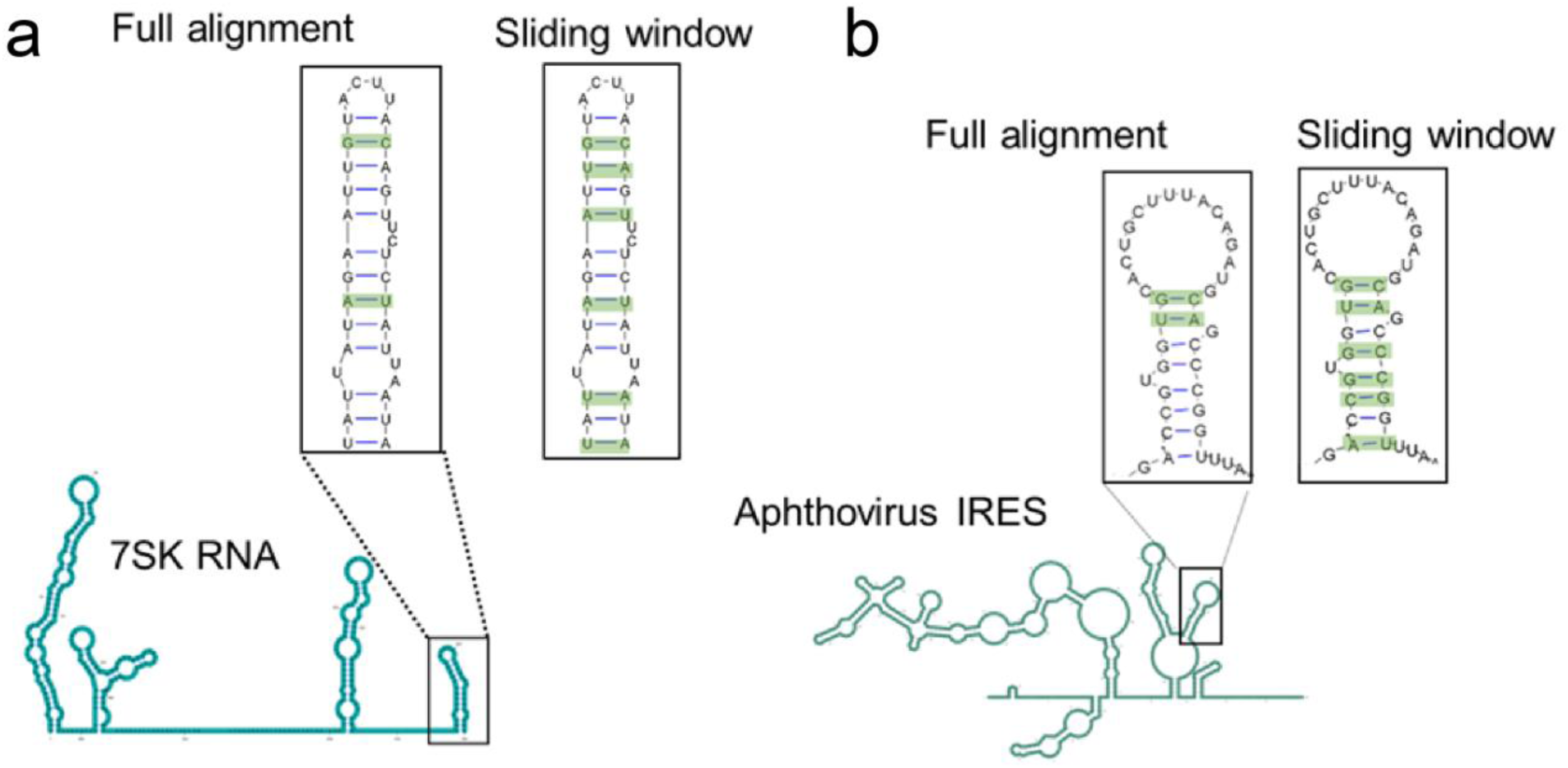
Sliding windows analysis improves R-scape performance on long alignments. In both model cases tested in this study, 7SK RNA (**a**) and Aphthovirus IRES (**b**), R-scape identified four additional base-pairs when the analysis was run in sliding windows. The consensus secondary structure of each RNA is shown in the cartoon form below, and insets above show the covariation predictions for specific domains. Predicted covariant base pairs are highlighted in green.

Taken together (Figures 1-3), these results suggest that one might be able to increase the signal-to-noise ratio for predicting lncRNA covariations by maximizing the number of sequences (increasing alignment depth) and running R-scape analysis in short windows. Here, we applied both conditions to analyze the RepA region of lncRNA Xist. In a previous study, R-scape identified no significant base pair covariation in RepA structure^11^. However, the input alignment in that study was limited to ten sequences, which was beneath an empirical threshold value (~40 sequences) suggested in the very same paper. We therefore reanalyzed RepA using a recent, experimentally determined secondary structure^10^ and we included significantly more sequences in the alignment. As expected, just by adding more sequences we were able to identify covariation in RepA, but it was limited to a single base pair. Interestingly, this base pair is located within the functionally important repeat-five region^19^. To further improve the signal-to-noise ratio, we ran R-scape on short (500-nt) overlapping windows, tiling the entire RNA. Using this procedure, R-scape identified five statistically significant covariant base pairs: two in domain I and three in domain II of the lncRNA RepA (Figure 4). Importantly, each of the covariant base pairs detected are in the regions of high structural confidence (low Shannon entropy, see Liu et al.,^10^ and Supplementary figure 2). It is also worth highlighting that the three base pairs in domain II identified by R-scape are in proximity to long-range crosslink sites identified by Liu et al. 2017^10^, and to a stretch of conserved base sequence, suggesting that even though R-scape identified only five base pairs, they are consistent with experimental studies and are likely to be functionally important.

**Figure 4.**
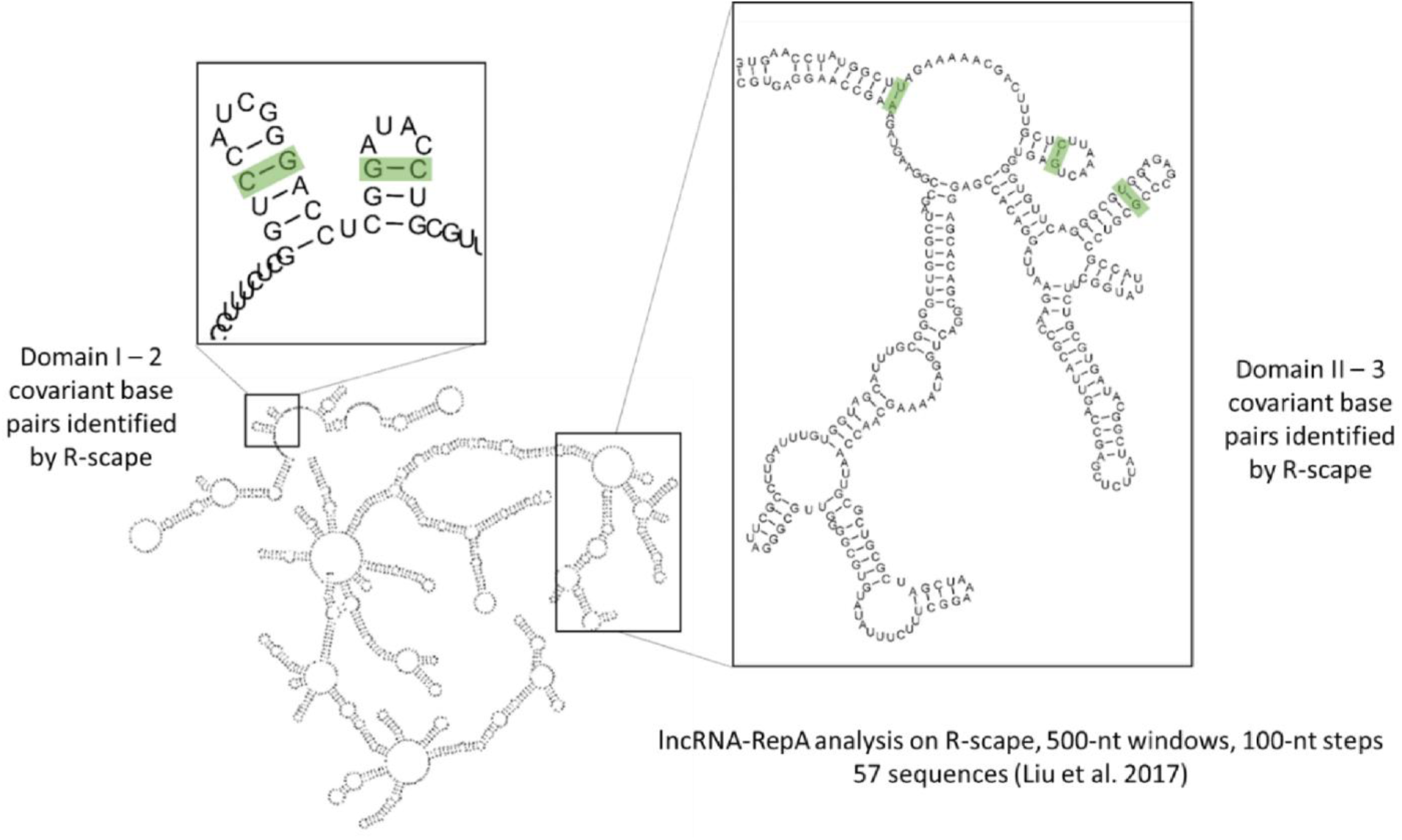
R-scape analysis on lncRNA RepA’s recently published structure (Liu et al. 2017). The use of an alignment containing 57 sequences was coupled with a sliding windows approach in order to improve covariation analysis on R-scape. The experimentally determined secondary structure of the lncRNA is represented in the figure with insets showing the covariant basepairs (green boxes) identified by R-scape on specific motifs of domain I and domain II of RepA (left and right insets, respectively).

Up to this point, our analysis suggests that the default parameters in R-scape are exceedingly stringent and that they may not be sufficiently sensitive to predict covariation with reduced alignment depth and low phylogenetic diversity, which are features inherent to most current lncRNA alignments (Xist, HOTAIR, SRA, etc). Most telling, R-scape failed to detect significant covariation when faced with similar alignments even for well-structured RNAs such as ribosomal RNAs, snRNAs and the eukaryotic ribozyme RNAseP, suggesting that more sequencing data is required to provide sufficient alignment depth for lncRNA structural conservation analysis on R-scape. Given the plethora of lncRNA genes and their implicated roles in human diseases, there is an urgent need for better tools and metrics to identify conserved structures and associated functions of these giant molecules.

We therefore asked whether other metrics could improve the performance of R-scape on long RNA molecules. RNAalifold with stacking^20^ (B^s^_i,j_ renamed in Rivas et al., as RAFS) was previously shown to be among the best performing covariation metric available and it has been extensively validated in several RNA structure prediction platforms, where it is frequently combined with structural stability metrics^21-23^. However, Rivas et al^11^ have argued that the G-test statistic (GT) performs better than RAFS in terms of positive predictive value (PPV) and thus would be less prone to false positive discovery. To get a sense of the tradeoff between sensitivity and PPV within these two metrics, we reanalyzed the original R-scape test set (104 RFAM alignments) with default parameters. We used average product correction (APC) as it was shown to improve the performance of both GT and RAFS (both renamed, then, as APC-GT and APC-RAFS)^11^. First, we measured the difference in sensitivity and PPV of these two metrics by varying the E-value threshold (Supplementary Figure 3). The PPV value for APC-RAFS gets worse than APC-GT (> 5%) only for relatively high E-values (> 0.1). However, at the default Evalue threshold of 0.05, APC-RAFS resulted in much higher sensitivity (~84%) relative to APC-GT (~64%), with a PPV compromise of less than 4%, suggesting that APC-RAFS is in fact a more robust metric than APC-GT.

Next, we tested the performance of these two metrics by varying the number of sequences in the input alignment (Supplementary Figure 3). Most strikingly, APC-RAFS achieved 63% sensitivity with only 20 sequences in the alignment compared to APC-GT, which resulted in only 40% sensitivity with the same input. We then used APC-RAFS to score the same alignments from Figure 1 (Supplementary Figs. 4 and 5) and observed a significant improvement in covariation detection under restricted conditions (fewer sequences, increased average pairwise identity and decreased phylogenetic diversity), relative to the original analysis using APC-GT. Remarkably, the eukaryotic RNAseP case showed a dramatic 45% sensitivity increase upon subsampled alignment analysis with APC-RAFS relative to APC-GT, and an even higher improvement (49%) on the mammalian sequence alignment. In all cases, the use of APC-RAFS on restricted alignments improved the overall covariation output when compared to APC-GT with no compromise to specificity as given by PPV (Supplementary Fig. 5), indicating that APC-RAFS is able to at least partly overcome the negative effects of lncRNA-like restrictions on R-scape predictive power while preserving statistical rigor. All these observations suggest that APC-RAFS is a highly robust metric for RNA covariation analysis with R-scape (null-model based analysis) and, most importantly, the most suitable method for alignments with the restrictions normally found in lncRNAs.

Based on the above observations, we utilized APC-RAFS to analyze the published structural alignments for full-length lncRNA-RepA and Domain I of lncRNA-HOTAIR^9^ (Fig. 5) and found that R-scape is now able to support covariation of numerous base pairs in both RNAs. We identified 16 covariant base pairs within the full-length lncRNA-RepA when the alignment was analyzed in overlapping 500-nt windows tiling the RNA every 100 nt. In this case, 9 out of 10 helical motifs with covariant base pairs flagged by R-scape/APC-RAFS were also suggested to be conserved in previous empirical studies^10^. Within HOTAIR domain I, 24 base pairs were flagged as covariant by R-scape/APC-RAFS in 10 helical segments of this region. Also, in this case, most helices where APC-RAFS found covariant base pairs overlapped with helices previously suggested as structurally conserved in domain I of HOTAIR^9^. These results strongly suggest that APC-RAFS can be used within R-scape to improve covariation analysis of lncRNA structure, confirming the conclusions from previous studies and highlighting the presence of conserved structured regions in lncRNAs HOTAIR and RepA.

**Figure 5.**
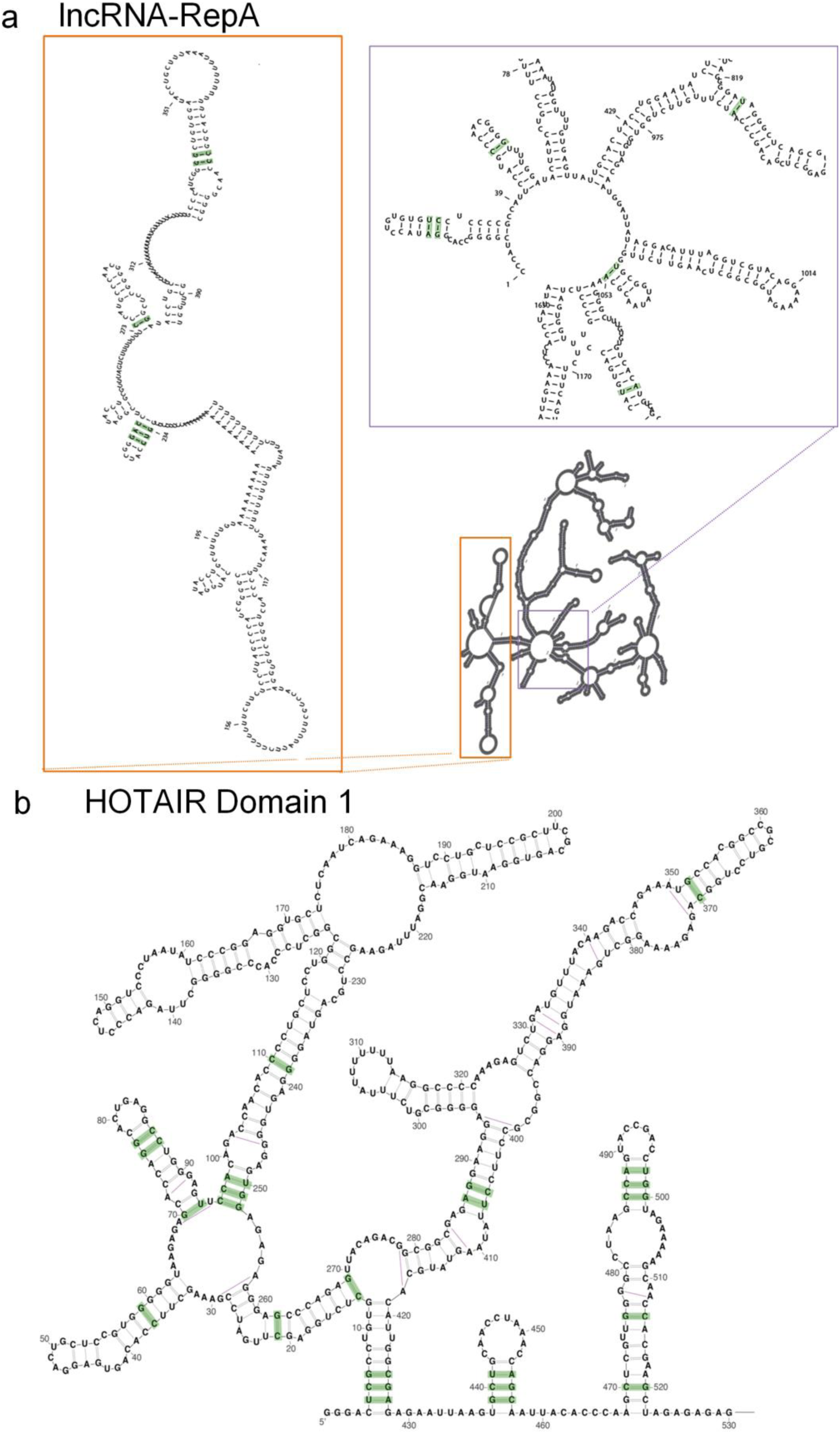
R-scape analysis on lncRNAs RepA and HOTAIR using APC-RAFS as the covariation metric. (**a**) The experimental secondary structure map of full-length lncRNA RepA is shown and covariant basepairs identified on specific motifs by R-scape using APC-RAFS are indicated in green boxes. (**b**) The experimental secondary structure of domain I of HOTAIR is represented in the figure and the covariant basepairs identified by R-scape using APC-RAFS are shown in green boxes.

In conclusion, we show that R-scape default parameters are not applicable to lncRNAs, but that R-scape is capable of identifying covariation when appropriately parameterized. We suggest that increased alignment depth, sliding windows approach and a more sensitive statistical metric, the APC-RAFS, are parameters that may help R-scape to identify conserved structural elements in large molecules such as lncRNAs. By combining these approaches, we were able to detect significant covarying base pairs in the experimental structures of lncRNAs HOTAIR and RepA. We hope that the results and approaches reported here provide improved tools for meeting the challenges inherent to studying lncRNA molecules and that they facilitate future studies and method development.

## Methods

### R-scape analysis

Seed alignments for tRNA, 5S ribosomal RNA, 5.8S ribosomal RNA, eukaryotic RNase P, U2 snRNA, U5 snRNA, small subunit ribosomal RNA (SS rRNA), 7SK, Aphthovirus IRES and SAM-I Riboswitch were downloaded from the RFAM database (RFAM v13.0). To obtain alignments restricted to mammals, mammalian sequences were manually extracted from each RNA family in the Rfam database and then aligned using Infernal (version 1.1.2). Sub-sampling analysis was performed by randomly selecting sequences using the ‘submsa’ option. The average pairwise identity (figure 1) was controlled using the ‘maxid’ option. The parameters and RFAM family IDs for all original and derived alignments are listed in Supplementary Figure 1. All analyses using R-scape were carried out at the default E-value (0.05), unless otherwise specified in the text.

Sliding window analyses were carried out with the “window” and “slide” options on R-scape, to define window size and sliding step of the R-scape search, respectively. Window size was varied between 50 - 500 nt, depending on the RNA length and structure, thereby ensuring that intact helices could be contained within the chosen window size.

The original R-scape test set was downloaded from the Eddy lab website (http://eddylab.org/R-scape/). The average product corrected RNAalifold with stacking (APC-RAFS) and G-test (APC-GT) statistics were compared using the “RAFSp” and “GTp” options respectively, by varying E-value thresholds and the number of sequences in the alignment.

## Acknowledgments

We thank Dr. Thayne Dickey for thoughtful comments on the manuscript. S.S. is supported by start-up funds from Drexel University College of Medicine and a CURE grant from the Pennsylvania Department of Health. R.C.A.T. was supported by NIH Grant RO1 50313. A.M.P. is an investigator with the Howard Hughes Medical Institute.

## Author contributions

S.S., R.C.A.T and A.M.P designed research and wrote the manuscript. S.S. and R.C.A.T carried out the experiments.

## Supplementary figure legends

**Supplementary Figure 1.**
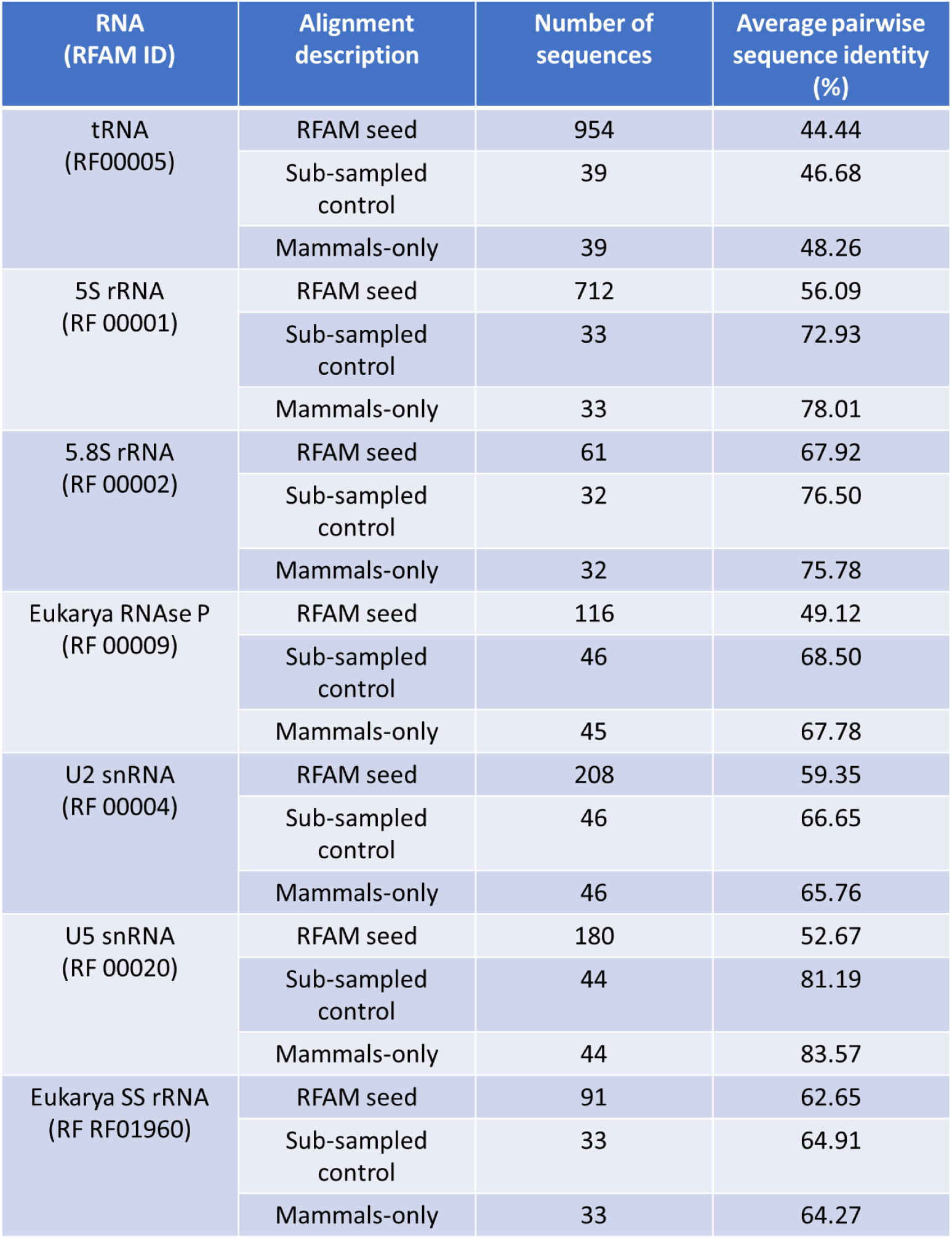
Parameters of structural alignments used for the R-scape analysis presented in Figure 1. RFAM IDs are indicated for each RNA family and the number of sequences and average pairwise sequence identity of each individual alignment (seed alignment, sub-sampled and mammalian sequences) are listed.

**Supplementary Figure 2.**
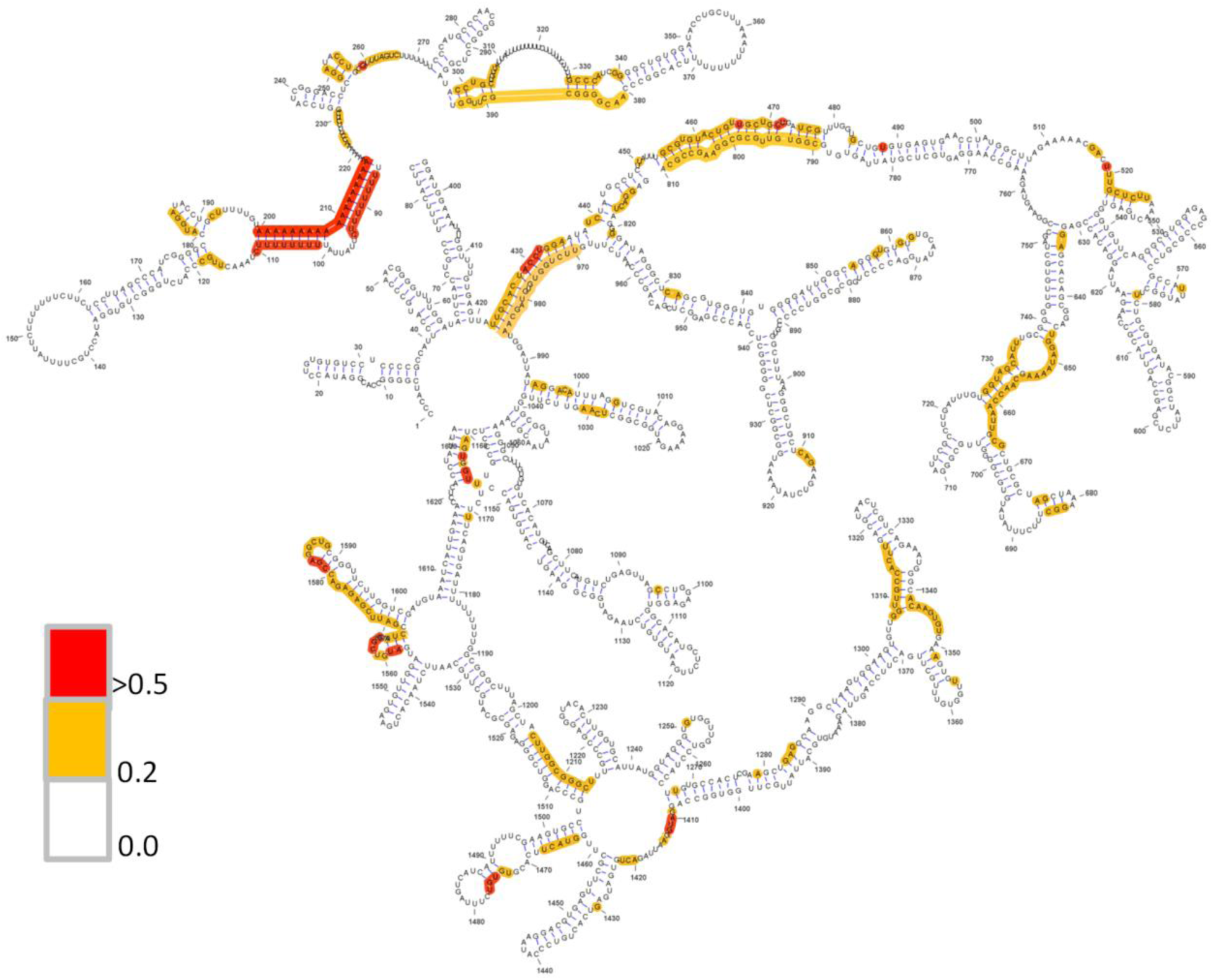
Shannon entropy values mapped onto the experimental secondary structure map of lncRNA-RepA (Adapted from Liu et al. 2017). Nucleotides with high Shannon entropy values are represented in red (> 0.5) circles; those with medium values (0.2-0.5) are represented in yellow circles. Nucleotides with low Shannon entropy (< 0.2) are not highlighted in the map.

**Supplementary Figure 3.**
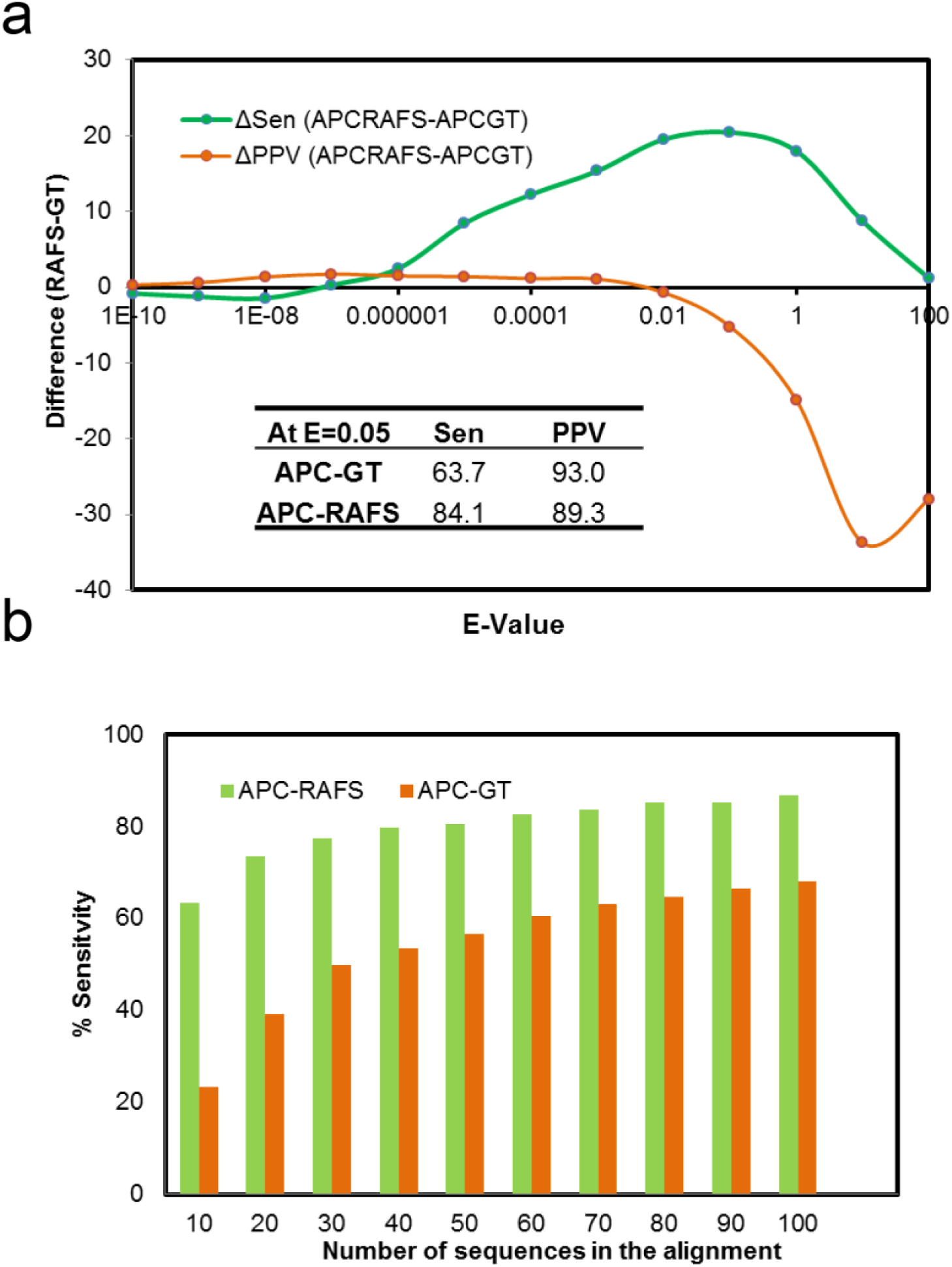
Comparison between the APC-GT and APC-RAFS covariation statistics currently implemented in R-scape. (**a**) Difference in Sensitivity (Sen) and Positive Predictive Value (PPV) between APC-GT and APC-RAFS at various E-value thresholds. At the R-scape default E-value, APC-RAFS shows much better sensitivity over APC-GT. (**b**) Same analysis as in (**a**), now varying the number of sequences in the alignments to include a range more commonly found in lncRNA alignments. At a fixed E-value threshold (0.05), APC-RAFS results in superior sensitivity even with 10 sequences in the alignment.

**Supplementary Figure 4.**
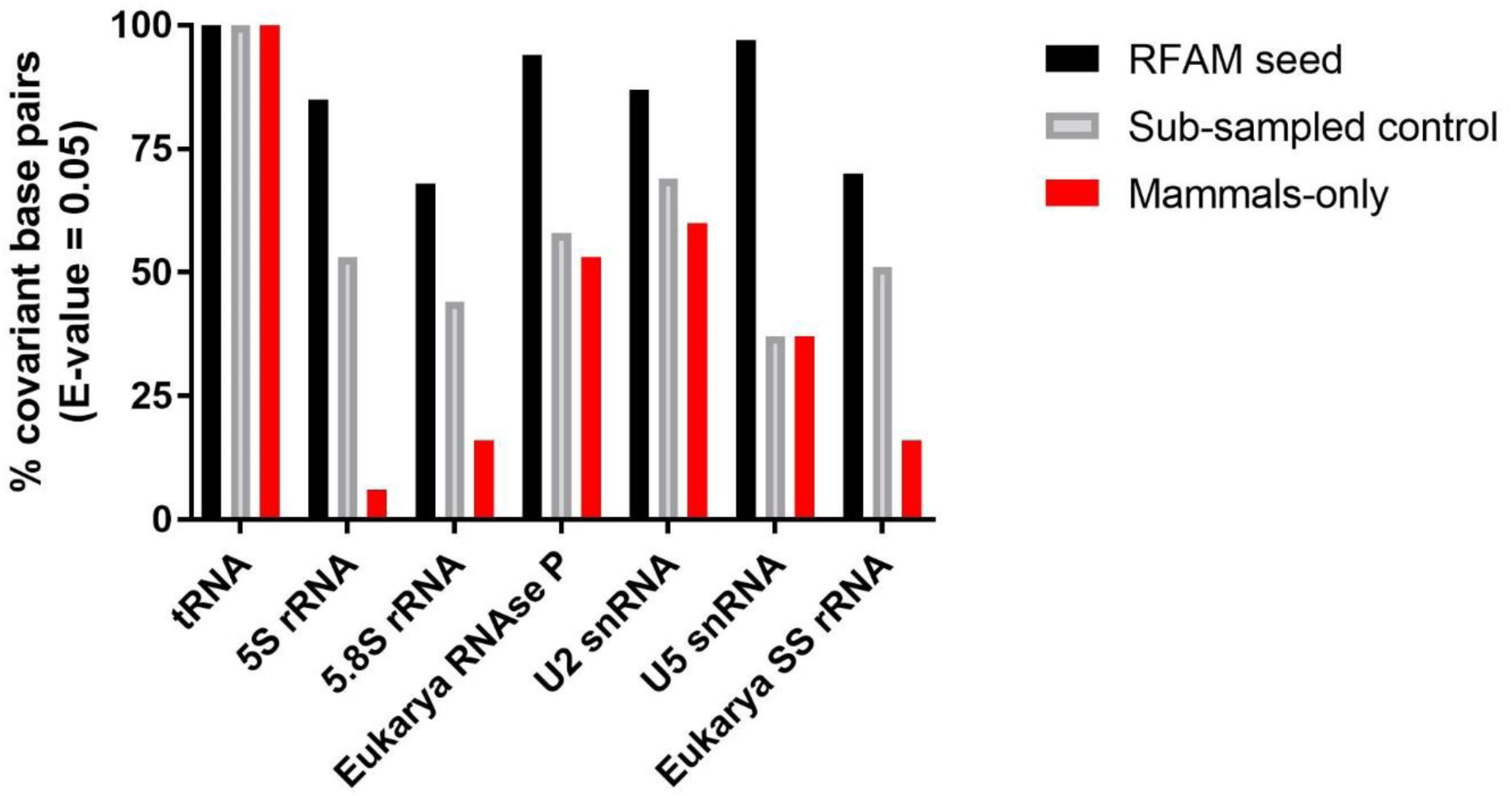
R-scape analysis using APC-RAFS as the covariation method on the same alignments used in Figure 1 of the main text. The percentage of covariant basepairs flagged by R-scape as statistically significant is shown in the graph for each tested RNA alignment.

**Supplementary Figure 5.**
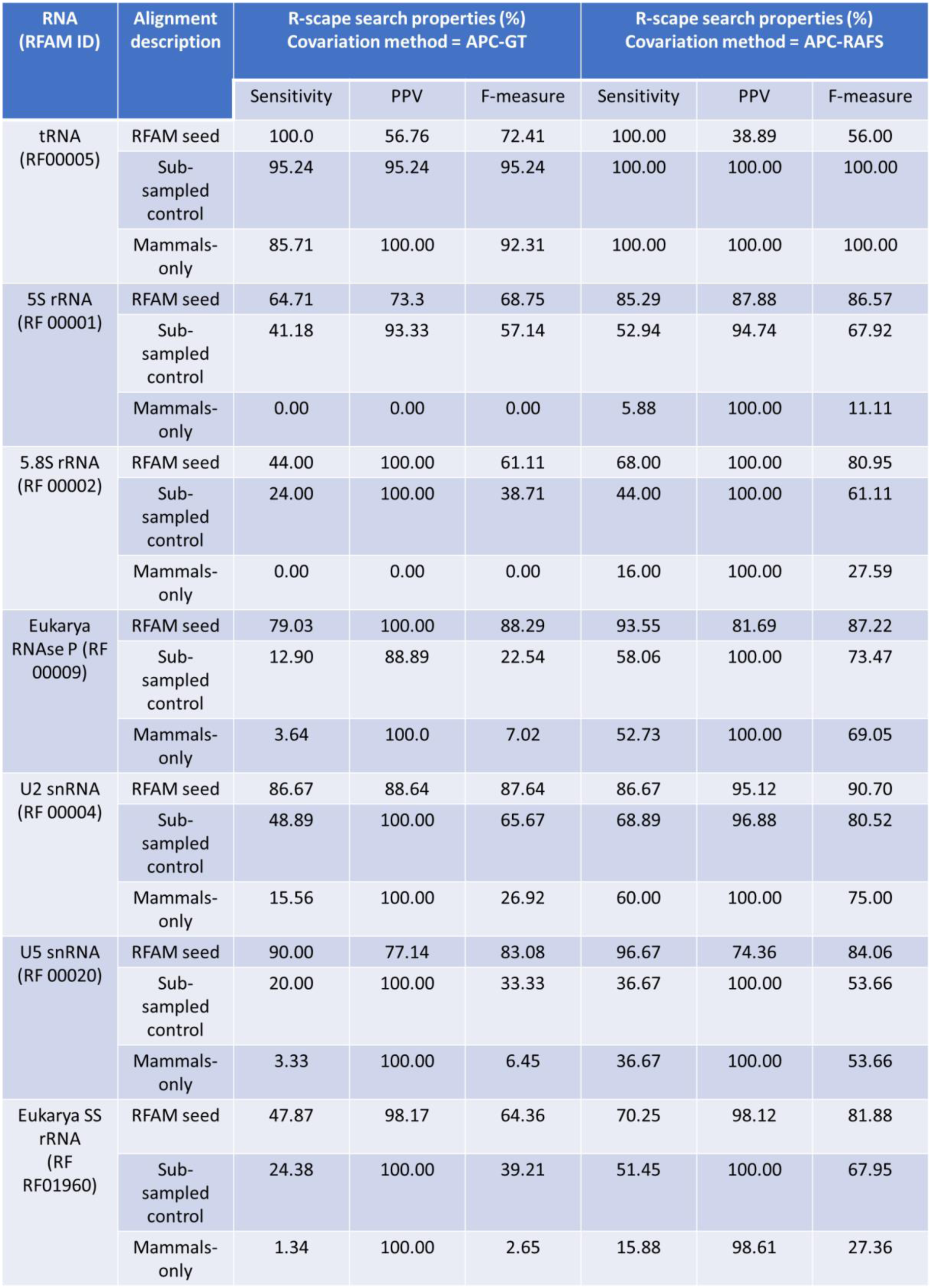
R-scape search parameters for the alignments referred to in Figure 1 and Supplementary Figure 1, comparing APC-GT and APC-RAFS covariation statistics. RFAM IDs are indicated for each RNA family. The percent values of sensitivity, positive predictive value and F-measure of each R-scape search were obtained from the analysis output for each alignment (seed alignment, sub-sampled and mammalian sequences) using both covariation methods, APC-GT (R-scape’s default) and APC-RAFS.

## References

1. Schmitt, A.M. & Chang, H.Y. Long Noncoding RNAs: At the Intersection of Cancer and Chromatin Biology. Cold Spring Harbor Perspectives in Medicine 7 (2017).

2. Pyle, Anna M. Looking at LncRNAs with the Ribozyme Toolkit. Molecular Cell 56, 13–17 (2014).

3. Mustoe, A.M. et al. Pervasive Regulatory Functions of mRNA Structure Revealed by High-Resolution SHAPE Probing. Cell 173, 181–195 e118 (2018).

4. Pirakitikulr, N. et al. The Coding Region of the HCV Genome Contains a Network of Regulatory RNA Structures. Mol Cell 62, 111–120 (2016).

5. Fu, Y. et al. Discovery of Novel ncRNA Sequences in Multiple Genome Alignments on the Basis of Conserved and Stable Secondary Structures. PloS one 10, e0130200 (2015).

6. Will, S. et al. LocARNA-P: accurate boundary prediction and improved detection of structural RNAs. Rna 18, 900–914 (2012).

7. Gruber, A.R. et al. RNAz 2.0: improved noncoding RNA detection. Pacific Symposium on Biocomputing. Pacific Symposium on Biocomputing, 69–79 (2010).

8. Novikova, I.V., Hennelly, S.P. & Sanbonmatsu, K.Y. Structural architecture of the human long non-coding RNA, steroid receptor RNA activator. Nucleic Acids Research 40, 5034–5051 (2012).

9. Somarowthu, S. et al. HOTAIR Forms an Intricate and Modular Secondary Structure. Molecular Cell 58, 353–361 (2015).

10. Liu, F., Somarowthu, S. & Pyle, A.M. Visualizing the secondary and tertiary architectural domains of lncRNA RepA. Nature Chemical Biology 13, 282 (2017).

11. Rivas, E., Clements, J. & Eddy, S.R. A statistical test for conserved RNA structure shows lack of evidence for structure in lncRNAs. Nature Methods 14, 45 (2016).

12. Nitsche, A. & Stadler, P.F. Evolutionary clues in lncRNAs. Wiley Interdisciplinary Reviews: RNA 8, e1376 (2017).

13. Kalvari, I. et al. Rfam 13.0: shifting to a genome-centric resource for non-coding RNA families. Nucleic Acids Research 46, D335–D342 (2018).

14. Smith, M.A. et al. Widespread purifying selection on RNA structure in mammals. Nucleic Acids Research 41, 8220–8236 (2013).

15. Hezroni, H. et al. Principles of long noncoding RNA evolution derived from direct comparison of transcriptomes in 17 species. Cell Rep 11, 1110–1122 (2015).

16. Chillon, I. & Pyle, A.M. Inverted repeat Alu elements in the human lincRNA-p21 adopt a conserved secondary structure that regulates RNA function. Nucleic Acids Res 44, 9462–9471 (2016).

17. Kuraku, S. et al. aLeaves facilitates on-demand exploration of metazoan gene family trees on MAFFT sequence alignment server with enhanced interactivity. Nucleic Acids Research 41, W22–W28 (2013).

18. Palazzo, A.F. & Lee, E.S. Non-coding RNA: what is functional and what is junk? Frontiers in Genetics 6 (2015).

19. Pintacuda, G., Young, A.N. & Cerase, A. Function by Structure: Spotlights on Xist Long Non-coding RNA. Frontiers in Molecular Biosciences 4, 90 (2017).

20. Lindgreen, S., Gardner, P.P. & Krogh, A. Measuring covariation in RNA alignments: physical realism improves information measures. Bioinformatics 22, 2988–2995 (2006).

21. Washietl, S., Hofacker, I.L. & Stadler, P.F. Fast and reliable prediction of noncoding RNAs. Proceedings of the National Academy of Sciences of the United States of America 102, 2454–459 (2005).

22. Hofacker, I.L. RNA consensus structure prediction with RNAalifold. Methods in molecular biology 395, 527–544 (2007).

23. Bernhart, S.H. et al. RNAalifold: improved consensus structure prediction for RNA alignments. BMC bioinformatics 9, 474 (2008).

